# Universal principles of protein resource allocation among cellular organelles revealed by yeast large-scale absolute quantitative proteomics

**DOI:** 10.1101/2025.09.04.674116

**Authors:** Hongzhong Lu, Zhendong Zhang, Siwei He, Yucan Zhang, Yuguang Wang, Fei Tao, Jens Nielsen

## Abstract

The presence of organelles is a hallmark distinguishing eukarya from bacteria and archaea, and this culminates in compartmentalization of cellular metabolism and subsequent metabolic specialization. Here we established a dataset encompassing over 300 absolute quantitative proteomes, the largest to date, across two yeast species under diverse experimental conditions. Leveraging big data analysis, formula fitting, and machine learning models, quantitative correlations among protein abundance, organelle-level resource distribution, and cellular phenotypes were elucidated at a system level. We found that protein resources always exhibit robust and precise distribution at the organelle level across distinct conditions. Specifically, at high specific growth rates, the protein mass fraction from some main organelles, i.e., peroxisome and nucleus, is consistently reduced to offset the increasing protein resource demand from the ribosome. Meanwhile, we found that the nutrition limitation could induce resource recycling by upregulating protein resources within the vacuole and lipid droplets to sustain stress adaptation. Importantly, our integrative analysis demonstrates that protein mass fraction from less than 4 organelles (e.g., nucleus and ribosomes) can accurately predict diverse yeast physiological parameters (e.g., specific growth rate, oxygen uptake rate), and a core set of 37 proteins could predict resource allocation among 24 main organelles and sub-organelles with high accuracy (average R^2 > 0.9). Finally, we found organelle resource allocation reflects the divergence of yeast species. For example, anaerobic conditions and respiratory suppression have less influence on Crabtree-positive yeast, i.e., *Saccharomyces cerevisiae*, with respect to organelle resource allocation but have a larger effect on the Crabtree-negative yeast *Issatchenkia orientalis*, thus suggesting that cellular resources have facilitated adaptive evolution. In summary, the high-quality, genome-scale quantitative proteomic dataset for yeast species offers an unprecedented opportunity for understanding the basic principles underlying resource allocation at the organelle level, laying theoretical foundations for precision engineering of cell factories in synthetic biology. The resource used in this study is available at https://yeast-proteome-database.streamlit.app/.

## Introduction

The evolutionary emergence of organelles and their associated membranes marked a pivotal transition in the development of eukaryal cells. This enabled metabolic specialization and spatial partitioning of biochemical pathways (1). Compartmentalization allows cells to optimize resource utilization, mitigate biochemical incompatibilities, and improve the catalytic efficiency of enzymes through the spatial proximity effect (2). Particularly, key organelles, like mitochondria, peroxisomes, and the nucleus, play significant roles in various metabolic activities, such as energy production, redox balance, genetic regulation, protein synthesis and degradation, et al. The division of labour and inter-organellar crosstalk not only sustain fundamental cellular processes but also establish a hierarchically organized metabolic network, forming the foundation for advanced biological functions. However, despite their central role in eukaryal physiology and growing significance for metabolic engineering (3), fundamental principles governing how cells allocate protein resources among organelles is still not well understood.

Recent advances in quantitative proteome analysis based on mass spectrometry (4) have pioneered systems biology studies of *Saccharomyces cerevisiae*, a key model for eukaryal biology. Quantitative proteome analysis has enabled profiling the dynamic changes of protein abundance in yeast under varying conditions, i.e., nutrition limitation (5, 6), growth phase shifting (7), gene perturbation (8) and cell cycle switching (9, 10). The subsequent comparative analysis of proteomics between distinct conditions helps to illustrate how cell metabolism is controlled to adapt to various perturbations (11). It clearly shows that the abundances of some enzymes increases linearly with specific growth rate (12), the so-called growth laws. It is also found that the synthesis of enzymes in some organelles or pathways is more than needed, which is related to protein resource pre-allocation for more flexible adaptation under fluctuating conditions (13, 14). Moreover, the comparative proteomic analysis between Crabtree-positive and -negative yeast species displayed that the two types of species are significantly different in efficiency of translation and energy generation (15, 16). Integrating proteomics with genomics and transcriptomics also provides valuable insights into how genetic and transcriptional regulation influences protein abundances (17). Thus, proteome analysis facilitates the in-depth studies of yeast metabolism with high resolution.

Protein location always reflect the proteins unique function in cellular metabolism as a result of evolution. Due to the importance of protein locations, various procedures have been developed to map the protein onto specific locations within the cell (18–22). Meanwhile, the progresses in absolute quantitative proteomic analysis provide unique chances for studying the variation in protein abundance at the organelle level, which helps to explore how resource allocation at the organelle level determines cell metabolic rewiring. As a example, mitochondrial proteome analysis by Baryshnikova et al. (7) quantified the organelle’s adaptive responses during diauxic shift, showcasing the protein resource reallocation from the cytosol to the mitochondria when growth switches from fermentative to respiratory metabolism. Proteome analysis at the organelle level enables the setting of new constraints for the metabolic model for the better simulation of cellular behavior. For example, the mitochondrial protein mass fraction calculated by absolute proteomics could be used to constrain metabolic models to simulate the Crabtree effect in yeast under at high specific growth rates (23). However, the current organelle-level proteome analysis only focuses on several intensively studied organelles, i.e., mitochondria, and the protein resource allocation at major organelles and sub-organelle levels was never investigated systematically across different species and growth conditions.

To address the above issues, we here reconstructed the largest standardized atlas of absolute quantitative proteomes for yeast, encompassing over 300 datasets spanning two yeast species – *S. cerevisiae* and *I. orientalis*. We integrated disparate proteomic datasets from chemostats, batch cultures, and stress conditions into a unified large-scale dataset, enabling cross-condition meta-analysis. Further, the protein resource allocation at organelle levels was thoroughly explored under different specific growth rates, stressful conditions, and yeast species to reveal the design principles of living systems. With the large-scale proteomic dataset, the precise quantitative links between protein abundance, organelle resource distribution, and cellular phenotypes were also investigated to probe the mechanisms underlying the resource allocation at organelle level. Overall, this work illustrates how eukarya optimize resource allocation among organelles for metabolic reprogramming under fluctuating environments by combining the protein location annotation and large-scale absolute quantitative proteomics, thus providing a useful reference for establishing the direct connection between organelles and cell functions.

## Results

### Reconstruction of large-scale yeast absolute proteome datasets

Proteome analysis has been widely employed to probe the metabolic activities of yeast, even though these datasets differ in the format and are stored in disparate locations for each study. To investigate how organelle structure influenced protein resource allocation, we firstly collected all absolute quantitative yeast proteomics or relative ones from previous studies (5, 7, 9, 11–13, 15–17, 24–27) (23, 28). All collected relative proteomics were transformed into quantitative ones based on normalization and rescaling approaches (29, 30). Overall, nearly 300 quantitative proteomes across 150 distinct conditions were collected in this work mainly for two yeast species – *S. cerevisiae* and *I. orientalis.* Note that 275 of 300 proteomes were specific to *S. cerevisiae* and the rest to *I. orientalis*; hence, unless specifically mentioned, the analysis in this work was mainly based on data from studies of *S. cerevisiae*.

The quality of yeast proteomics varied across datasets. For instance, the number of proteins detected by proteomics could range from 1,500 to 5,000 in the original datasets we gathered. To enhance the reliability of the compiled dataset, only high-quality proteomes with over ∼2,000 proteins detected in MS/MS were retained (Fig.1a, Methods). Meanwhile, considering that the cell geometric information (i.e., cell size and surface area) was not known for most conditions, all types of protein abundance were transformed as mass fraction (g/g_total protein_) for the subsequent analysis (Methods).

**Figure 1.**
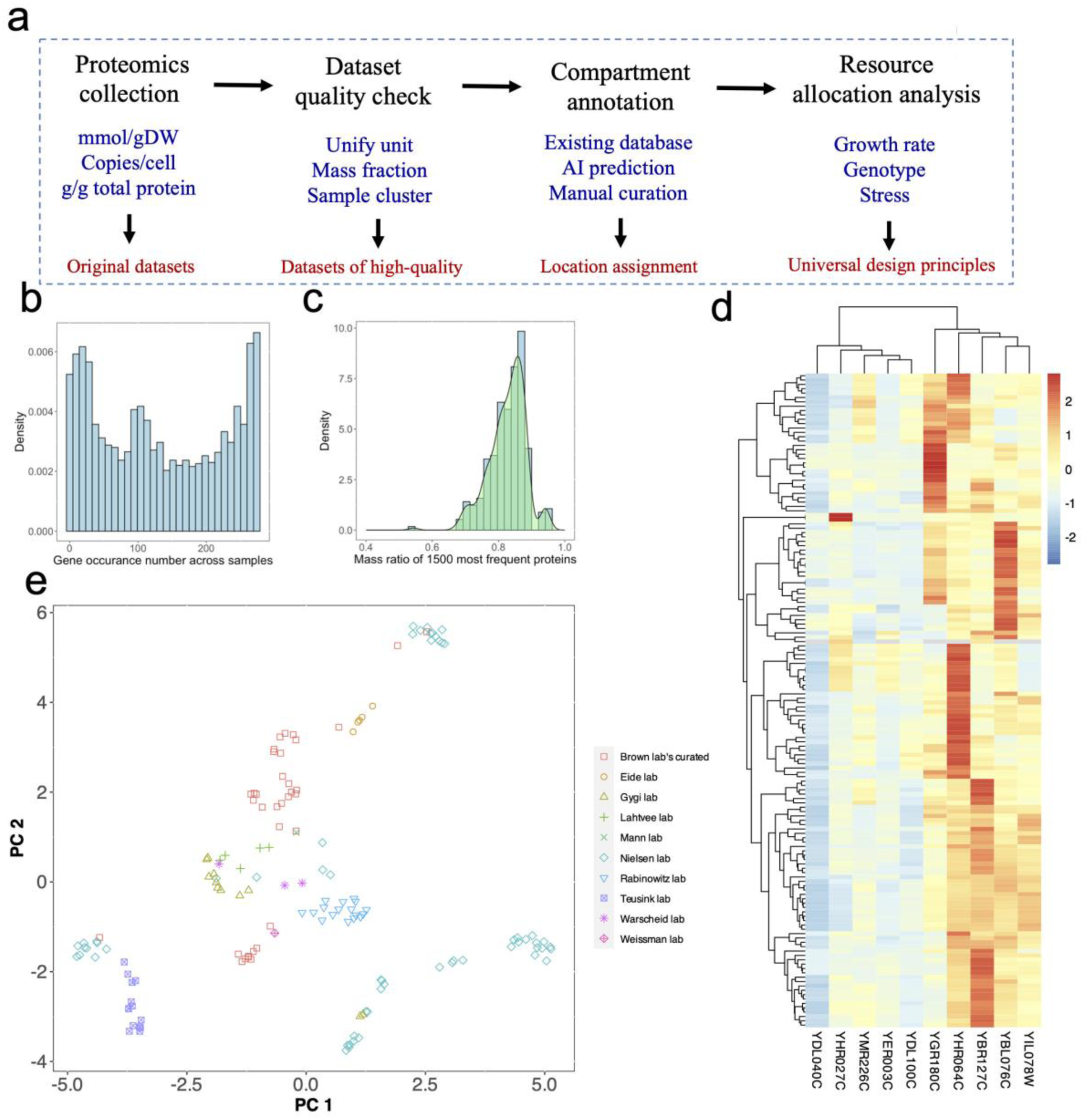
Collection and quality analysis of proteomics datasets for yeast. The whole pipeline of this study from the yeast proteomics collection to the distillation of novel biological insights (a). Protein occurrence number analysis across all samples collected by this study (b). Statistical analysis of the mass ratio of 1500 most frequently occurring proteins across samples (c). Conservation in mass fraction of proteins across 150 unique conditions. Here 10 different proteins were selected for the visualization (d). Clusters of samples based on the mass fraction of measured proteins which are not zero in over 50% of unique conditions across labs (e). Note that from each lab, there may exist multiple independent studies. For example, the proteomics from Nielsen lab encompassed 9 different studies.

### Yeast absolute proteomes exhibited consistency across samples

After quality analysis, we discovered that the yeast proteomes with unified mass fraction exhibited consistency, while some variances existed. To begin, the Pearson correlation coefficient (*ρ*) for any two samples was calculated, and it indicates that, in most cases, the *ρ* is over 0.5 (Fig.S1a), showcasing a relatively good consistency in **p**rotein **m**ass **f**raction (**PMF**) between any two samples. In comparison, it displays that the quantity of detected proteins varied significantly between samples from different labs or studies (Fig.S1b). In this regard, we examined the frequency of each protein detected in all samples and found some proteins were missing in nearly half of all samples (Fig.1b). One major reason may be due to the fact that these proteins are in extreme low abundance within cells. Overall, it shows that the accumulative mass fraction of the 1,500 most frequently measured proteins in all samples is over ∼0.7 (Fig.1c). It is also observed that, for each sample, the top 1,000 proteins with the highest abundance account for more than 90% of the mass of all measured proteins (Fig.S1c), demonstrating that proteome analysis from each study accurately captured the majority of the cellular protein pool. With PMF, it is convenient to assess the profile of protein abundance across all conditions. As shown in Fig.1d, it clearly hints that some proteins (i.e., YDL040C, YHR027C) have relatively lower abundance than other proteins (i.e., YBR127C, YHR064C), suggesting a consistent trend in protein expression across conditions. Further, the experimental conditions could be clustered according to the PMF. However, when all proteins in each sample were evaluated, we found that the samples from the same studies tend to cluster together (Fig.1e). Such a phenomenon can be caused by a variety of factors, including but not limited to batch effects, variation in strains and experimental conditions, different amounts of detected proteins in each study, et al. As a result, it should be given careful consideration when conducting meta-analyses of proteome datasets collected from different studies or labs.

By comparison, our dataset is nearly two folds larger than previous studies from (11, 31) (Fig.S1d) in the size of absolute quantitative proteomics. With those curated proteome datasets in hand, an online database (https://yeast-proteome-database.streamlit.app/) was created, allowing for the investigation and visualization of PMF trends across conditions. For example, using the online database, the correlation of log-transformed mass fractions for any two samples could be calculated and the PMF distribution across different samples could be displayed based on user preferences (Fig.S2). Thus, it is believed that this online database would be a useful resource for investigating the protein resource allocation across distinct growth environments. This online resource will be further expanded once additional yeast proteome datasets are available.

### Assigning organelles for each protein in yeast

Yeast possesses around 10 major organelles (32), each of which has unique metabolic function within the cell. Some major organelles could be further divided into sub-organelles. For example, as one single primary organelle, mitochondria could be further divided into Outer membrane, Intermembrane space, Inner membrane, Cristae and Matrix. To examine how protein resources are allocated at (sub-)organelle level, detailed cellular protein location annotation was performed. First, all the protein annotations were retrieved from the SGD database, which was then carefully curated using some recent literature reports (Method). As a result, almost all *S. cerevisiae* proteins were assigned to one or more organelles (or sub-organelles). For the latter, we calculated the mass fraction of proteins shared by two organelles per total protein mass using proteome data from unlimited growth conditions, and found that this mass fraction is smaller than 4% in most cases (Fig.S3a). As reported (10), some proteins could migrate dynamically into different organelles to execute the corresponding tasks in both organelles. To assess whether we can neglect these protein in our analysis, we recalculated the mass fraction of proteins which could transfer across two organelles under different cell cycles from (10) and found that the number is smaller than 1% in most cases (Fig.S3b), showcasing that this part of the proteome may not significantly influence our following analysis.

As shown in Fig.2a, the protein was unevenly distributed among different organelles. For example, some major organelles, i.e., nucleus and mitochondrion, contain over 1,000 proteins, while some smaller organelles, i.e., lipid droplet and P-body, only have less than 100 proteins. Compared to organelle annotation based on reliable experimental evidences, the number of proteins in some major organelles could be further refined (Fig.2b). Taking mitochondria and its sub-organelles as examples, according to a recent work (33), the protein list in each component of mitochondria was updated. As a result, the number of total proteins in mitochondria decreased from 1,193 to 1,127, while the number of proteins in the mitochondrial inner membrane (MIM) increased from 249 to 427 when compared to the protein annotation directly downloaded from the SGD database (34).

**Figure 2.**
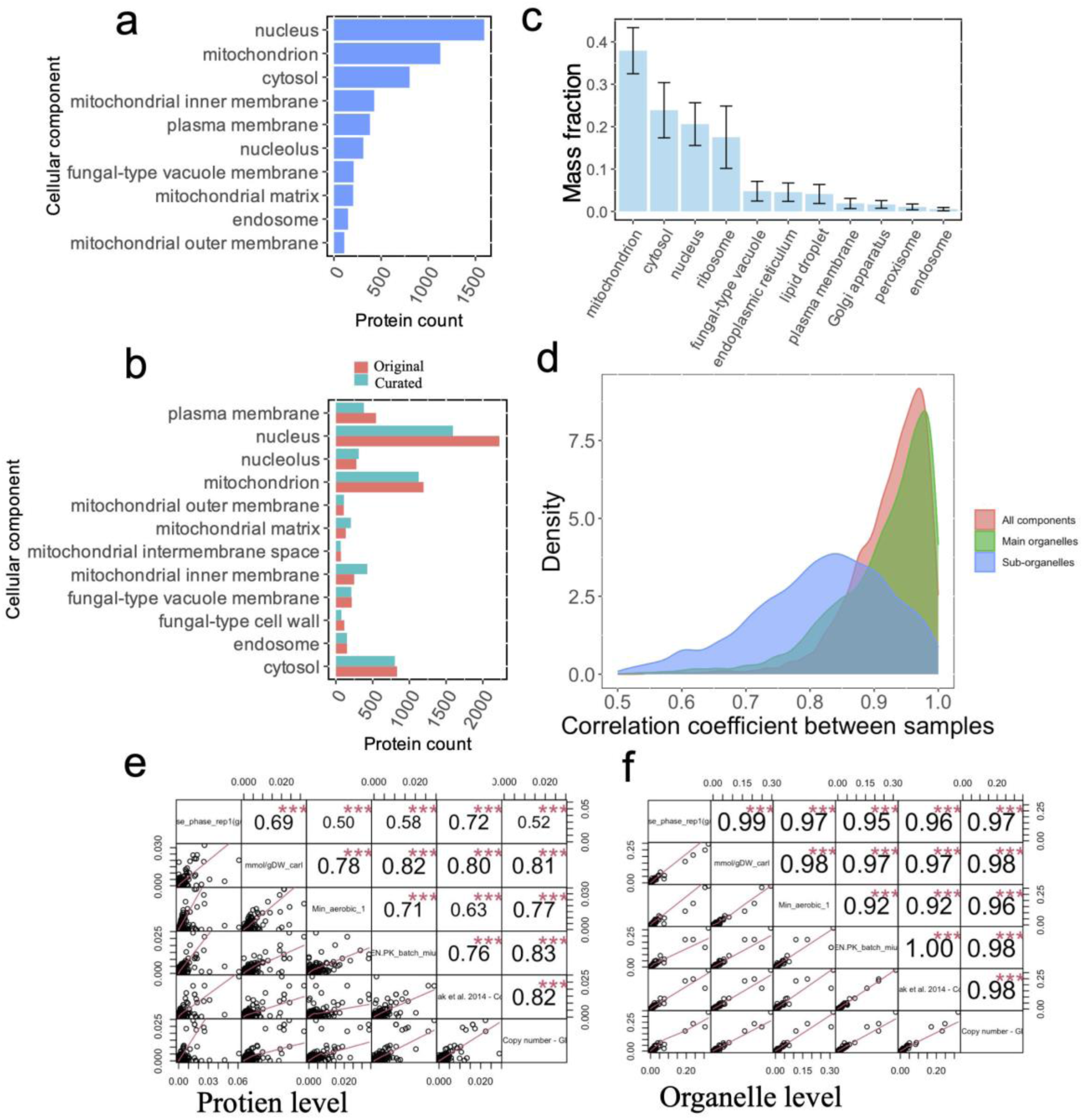
General trend in protein resource allocation at organelle level across nearly 150 unique conditions. Compartment annotation of *S. cerevisiae* proteins based on SGD database (a). Compartment curation based on experimental datasets (b). Conservation and variation in mass fraction for proteins from main organelles (c). Distribution of correlation coefficient calculated by protein mass fraction at organelle level across conditions (d). Pair-wise correlation coefficient calculated by mass fraction at protein level and at organelle level among 6 samples under batch cultivations from different studies or labs (e and f). ***: *P* value < 0.001.

### Robustness of resource allocations at organelle level

Combining PMF and protein locations, we computed the PMF for each organelle across 150 unique conditions to explore the resource allocation at the organelle and sub-organelle levels. As a whole, the protein resource is also unevenly distributed among different organelles. We found that, on average, the mitochondrion has the highest percentage in mass ratio of protein resource (>35%), followed by the cytosol (∼24%), nucleus (∼21%), and ribosome (∼19%). Except for the four major organelles, the mass fraction of protein for other major organelles was all smaller than 10% (Fig.2c).

Next, we found that there exist correlations between PMFs of organelles (or sub-organelles) across the different datasets. As some specific sub-organelle contain a few proteins, all organelles and sub-organelles were divided into three types: large components with PMF greater than 0.0035; medium components with PMF between 0.00035-0.0035; small components with PMF smaller than 0.00035 (Fig.S4). It is shown that protein resource allocation among the large components exhibits higher robustness as the corresponding correlation coefficient in organelle PMF is over 0.8 in most pairs of two samples (Fig.2d, Fig.S4a). By comparison, the correlation coefficient was decreased for medium and small components (Fig.S4b and S4c), probably indicating that the corresponding protein resource allocation was more flexibly redistributed. It should also be noted that the fluctuation in mass fraction of proteins from medium and small components could be larger, as part of the proteins was not measured. If considering all cellular components, including large components and small ones, the correlation coefficients are kept at a high level (Fig.2d), showcasing that the large components take up a higher weight in this kind of analysis. Taken together, it hints that there seem to exist scale effects in resource allocation at cellular component levels; that is, resource allocation tends to be more robust for cellular components with higher mass fraction.

To further assess the robustness in protein resource allocation at the organelle level, the *ρ* in mass fraction at the protein level and organelle level were computed, respectively, between samples under batch cultivations. Obviously, it shows the *ρ* is in the range of 0.52-0.83 for protein-level mass fraction under different conditions, smaller than that calculated for organelle-level mass fraction, which is in the range of 0.92-1.00. Moreover, the relatively higher correlation at both the protein level and organelle level also showcases the high quality of the datasets we collected in this study.

The above analysis mainly focused on the PMFs of organelles. However, the protein and organelle geometric size could also have an influence on the resource allocation. To this end, we systematically computed the volumes of protein from each organelle, as well as the sectional area of proteins from each membrane of organelles. Similar to PMF, we could calculate the protein volume fraction (PVF) of specific organelles per total protein volume, and it is found that there exists high linear correlation between PMF and PVF (Fig.S5a). Thus, the above conclusions based on PMF also hold based on PVF. The high correlation between PMF and PVF is mainly due to the fact that the calculated protein volume is directly determined by molecular weight (Fig.S5b).

### Growth regulates protein resources redistribution at (sub-) organelle level

Previously, it was found that the protein resource allocation could be dominantly regulated by the specific growth rate (the so-called growth law) (12, 35). Here, based on the unified PMF across diverse conditions, we checked how growth regulate the protein resource allocations at the cellular organelle level. For this proteomics from carbon-limited chemostat cultivation was employed (5). Firstly, we computed *ρ* between PMF from each component, including the major and sub-organelles, according to the proteomics under a series of growth rates from carbon-limited chemostat cultivation. As shown in Fig.3a, we found there exists a positive correlation between PMFs for some organelles, i.e., ribosome and mitochondria inner membrane (Fig.3b), and growth under carbon-limited condition. Meanwhile, the negative correlation between PMF and growth was also observed for lots of organelles, i.e., vacuole and lipid droplet. It initially indicates that there exists a trade-off in protein resource allocation at the organelle level along with cellular growth.

**Figure 3.**
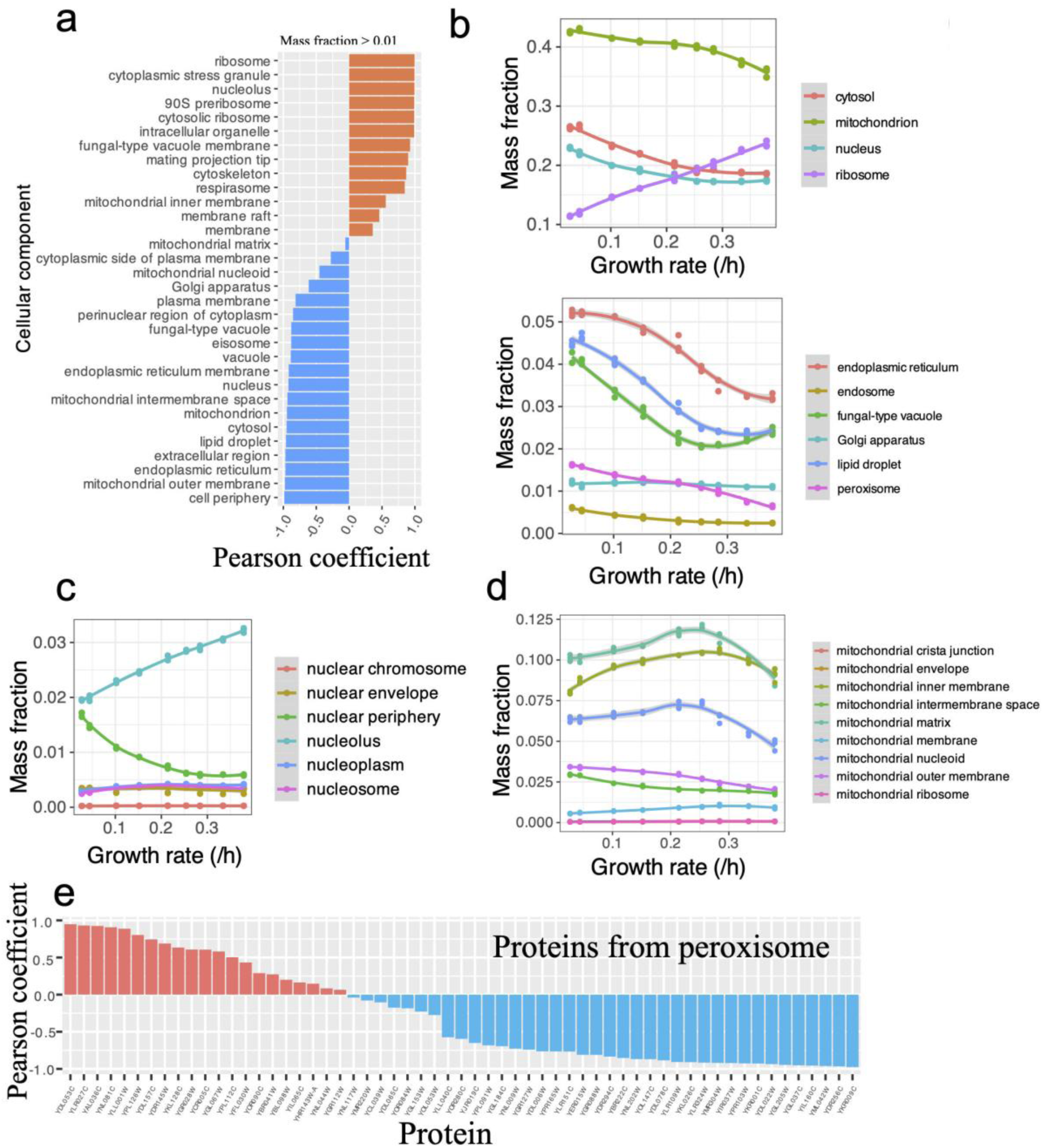
Growth rate significantly regulates protein resource redistribution at organelle level. Pearson coefficient between protein mass fraction at organelle level and the growth rate from carbon limited chemostat cultivation experiments (a). Detailed correlation between the main organelle’s protein mass fraction and the growth rate (b). Relative change in protein mass fraction for the sub-organelles from nucleolus along with the growth rate (c). Tendencies in changes of protein mass fraction for the sub-organelles from mitochondrion along with the growth rate (d). Pearson coefficient between mass fraction of each protein from peroxisome and the growth rate (e).

At the major organelle level, we found that only the mass fraction of ribosome increased consistently with growth rate under both nitrogen-and carbon-limited chemostat cultivations (Fig.3b, Fig.S6), while the mass fraction of proteins from other major organelles mostly decreased along with growth rate. It may be due to that the cell need to redistribute more resource into the protein synthesis machinery from other major organelles to meet the high demand for growth

Interestingly, if checking the resource allocation at the sub-organelle level from the nucleus, we found that the PMF of the nucleolus increases linearly with the cellular growth rate, although the PMF of the nucleus shows the opposite trend (Fig.3c, Fig.S6). Such a phenomenon may be due to the fact that the nucleolus within the nucleus is responsible for producing and assembling ribosomes. Thus, the higher demand in ribosome will correspondingly promote the increase in the PMF of the nucleolus. On the contrary, it shows that PMF of nuclear periphery decreased consistently as growth rate increased, showcasing the trade-off in resource allocation within nuclear. At a lower specific growth rates, the higher PMF of the nuclear periphery may represent a reserve of proteins, which is similar to the reserve of enzymes from metabolic super-pathways (13). It is speculated that as the PMF of the nucleus decreased at higher specific growth rates, the resource within the nucleus was reallocated from the nuclear periphery to the nucleolus to maintain the normal function of the nucleus.

The similar trade-off in resource allocations was also found for mitochondria, one key major organelle of eukaryal cells (Fig.3d). Different from the nucleus, it seems there exist constraints from mitochondria as a whole. We found the PMFs of four sub-organelles from mitochondria, including MIM and mitochondrial matrix, firstly increased with specific growth rate before the critical point and then decreased accordingly. It is well-known that *S. cerevisiae* has metabolic overflow under high specific growth rates. It is reported that once overflow started, more protein resources were channeled into the fermentation pathway from the respiration pathway; thus, the protein resources within mitochondria were withdrawn into the cytosol at the major organelle level. Here, our result, at the first time, clearly illustrates the tendencies in resource allocation at the major organelle level, as well as at the sub-organelle level of mitochondria, related to the occurrence of overflow.

The tendency in PMFs of the major organelle not only imposes constraint onto the changes in PMFs of its sub-organelles but also onto variation in mass fraction of each protein within the organelle. Taking peroxisome as an example, overall, the PMF of peroxisome decreased as growth increased (Fig.3b). Consequently, it shows that the PMFs for a larger ratio of proteins have a negative correlation with specific growth rate, which means cells tend to reduce the synthesis of proteins within the peroxisome to save the protein resource. By contrast, the PMF for some proteins within the peroxisome still increased, for example, YDL053C. However, when checking the detailed cellular location of those proteins with increased PMF, we found that they could be distributed within different organelles besides the peroxisome. Therefore, the different tendencies may potentially correlate to actual locations of the studied proteins. As the second example, the ρ between PMFs from nucleolus and growth was analysed (Fig.S7). Different from that of the peroxisome, the ρ for most proteins within the nucleolus is positive, consistent with the overall increased PMF of the nucleolus under higher specific growth rates. Despite this, some proteins, for example YNL007C, undergo a decrease in PMF, indicating the complex regulation in protein resource allocation within the nucleolus. Overall, the PMF was significantly affected by the total PMF of the organelles under different specific growth rates while the existing trade-offs could counter the influence from organelles.

Does the organelle itself really impose constraints on resource allocation? It is observed that a part of the proteome could be located in multiple organelles within a cell. It is doubted whether the above correlation between PMFs of organelles and specific growth rate was remarkable. Here, to answer this question, we randomly select hundreds of proteins (200, 500, and 1000), the number of which is in the range of the protein number within each major organelle, and then check the correlation between the PMFs of those randomly selected proteins and specific growth rate. Clearly, it shows that if randomly selected, not considering the protein location, there is no correlation between the PMF of sampled proteins and the specific growth rate, therefore showcasing that cellular compartment actually shapes the resource allocation within cells (Fig.S8).

### Quantitative correlations between protein abundance, organelle-level resource distribution and cellular phenotypes

From the aforementioned analysis, it indicates that there exists direct correlation between PMFs of organelles and cellular phenotypic datasets. Here we thoroughly explored how PMFs from key major organelles together could play a significant role in determining the cellular traits, not only specific growth rate, but also the specific uptake rates of glucose (q_glucose_) and oxygen (q_O2_) as well as the specific carbon dioxide production rate (q_CO2_). We firstly fitted the formulas using a larger proteomics dataset acquired under nitrogen-limited chemostat condition with diverse nitrogen sources and then tested the formulas using the above proteomics under carbon-limited chemostat condition (Fig.4a and 4b). In general, we found that, using the linear formula, less than 4 organelles’ PMFs could predict the cellular phenotypic datasets very well (Fig.S9). For the specific growth rate, only ribosome PMFs can predict it well, consistent with the previous reports that the growth rate is linearly correlated to the PMFs of ribosome (26). For q_glucose_ and q_CO2_, the PMFs of ribosome and nucleus could predict them better than only using one of them. By contrast, for q_O2_, four organelles’ PMF could also enhance the prediction performance. Overall, it could be concluded that organelle’s resource allocation is directly correlated to the cellular physiological parameters.

Thus, changes of PMFs from several key organelles could, to large extent, determine growth-related parameters of yeast, but what is the mechanism underlying the resource reallocation among different organelles? Can we find a group of proteins to predict the general resource allocation at organelle levels? To evaluate this, we built a model between PMF of all proteins and PMF of major organelles and sub-organelles from mitochondria and nuclear (Fig.S10) using a linear ridge regression model, in which all the proteomic datasets collected for *S. cerevisiae* were utilized. Through the feature importance analysis of ridge regression model, it shows that only 37 proteins could be used to predict the resource allocation among the major organelles (average R^2^ > 0.9, Fig.4c and Fig.S10), indicating organelle level resource allocation could be explained well by this minimal protein set. When doing KEGG pathway enrichment analysis (Fig. 4d), we found that these proteins could be significantly enriched in carbon metabolism (*P* value = 0.008), glycolysis pathway (*P* value = 0.017) and biosynthesis of secondary metabolites (*P* value = 0.017). We further probe the interaction of those 37 proteins based on the STRING database (36). As shown in Fig. 4e, the protein-protein interaction (PPI) network is highly interconnected, with average node degree =14.3 and PPI enrichment *P* value =2.04e-10, indicating that the density of interactions in the network is not randomly generated. The PPI network was mainly composed of three modules, 17 proteins in red are related to aerobic respiration, 8 proteins in blue are related to glycolysis/gluconeogenesis, and the green proteins are related to other pathways. PPI network analysis may hint that PMF of proteins from aerobic respiration and glycolysis/gluconeogenesis could be as meaningful indicators to reflect the general resource allocation at organelle level.

**Figure 4.**
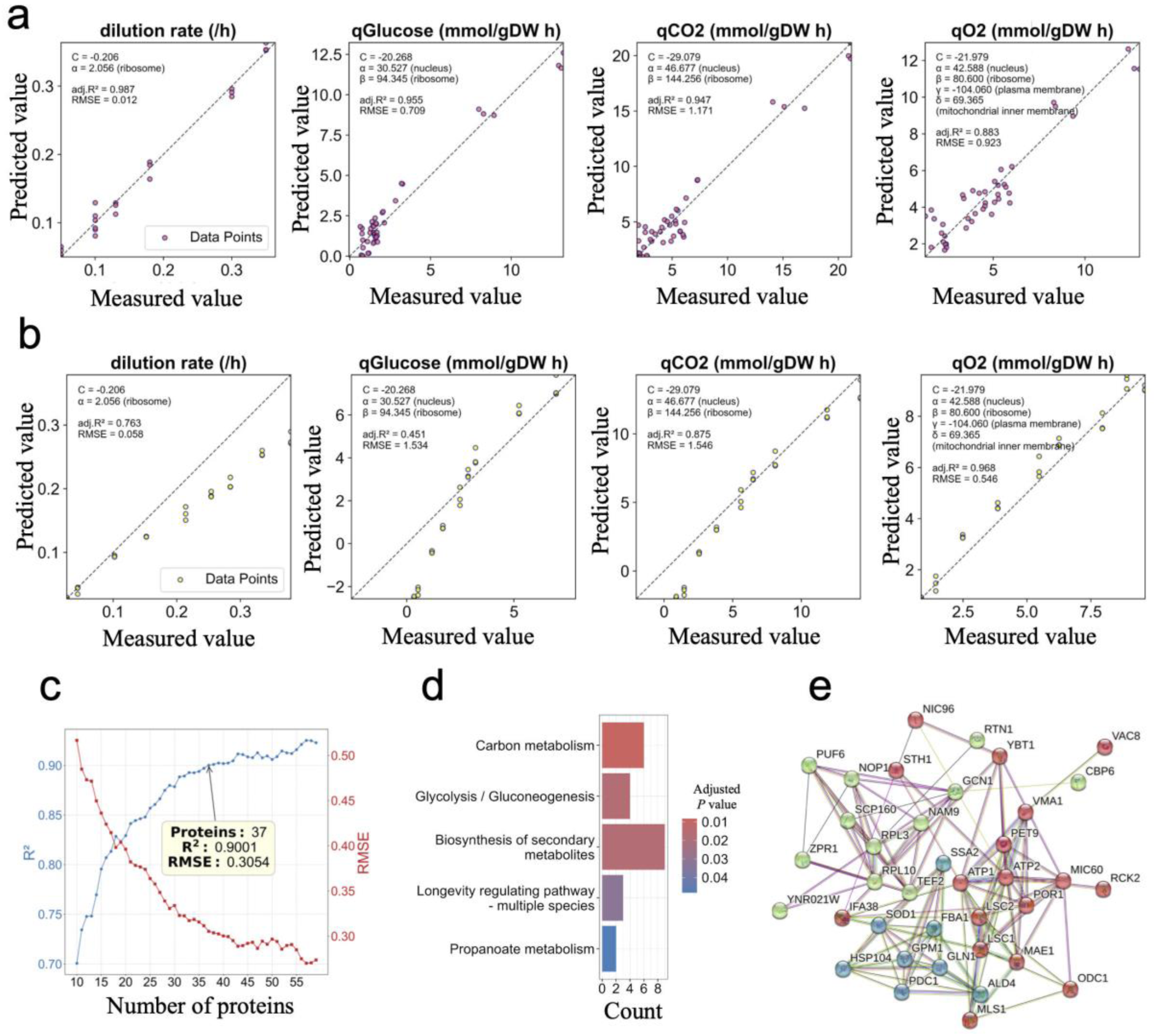
Quantitative correlations between protein abundance, organelle-level resource distribution and cellular phenotypes. Fitting of the linear correlations between the PMFs of organelles / sub-organelle and yeast physiological parameters using Yu’s large-scale dataset to obtain the formulas (a). Validation of the above formulas using Xia’s dataset (b). Relationship between the number of proteins used as input and the value of R^2^ ( or RMSE) when using the ridge regression model to predict the organelle level resource allocation from the mass fraction of proteins. To achieve R^2^ over 0.9, the minimal number of proteins is 37 (c). KEGG pathway enrichment for the 37 key proteins which could characterize the organel-level resource allocation (d). The PPI of the above 37 proteins obtained from string database with the average node degree =14.3 and the average local clustering coefficient =0.652. The 17 genes in red are related to aerobic respiration, the 8 genes in blue are related to glycolysis/gluconeogenesis, and the green genes are related to other pathways (e).

### Metabolic shift induced by nutrition limitation results in protein resource redistribution at organelle level

Yeast always encounters resource deficiencies in the environment, which may be accompanied with a metabolic shift to optimize resource utilization. It is known that nitrogen limitation could influence the cellular metabolism significantly. It is unclear how cellular protein resources are reallocated under different levels of nitrogen limitation. To alleviate the effects of regulation by growth rates, the proteomic dataset under different C/N ratios with growth rate controlled at 0.2h^-1^ using chemostat cultivation were reanalyzed in this study (13). In general, we found that the protein resource allocation at the organelle level exhibited high robustness, as the adjusted R^2^ between PMFs of organelles from different C/N ratios is over 0.94, though it is slightly decreased from 0.965 at C/N=30 to 0.941 at C/N=115 (Fig.5a). With the intensification of nitrogen resource limitation, we also found that the PMF of the ribosome decreased slightly. Note that the specific growth rate under different C/N ratios is kept at 0.2h^-1^, thus maybe some non-essential proteins from the ribosome were more degraded to meet the protein demand in other organelles. Interestingly, it is found that the PMFs for organelles responsible for degradation, like vacuole, are increased, which may accelerate the resource recycle under extreme N limitation (Fig. S11a).

**Figure 5.**
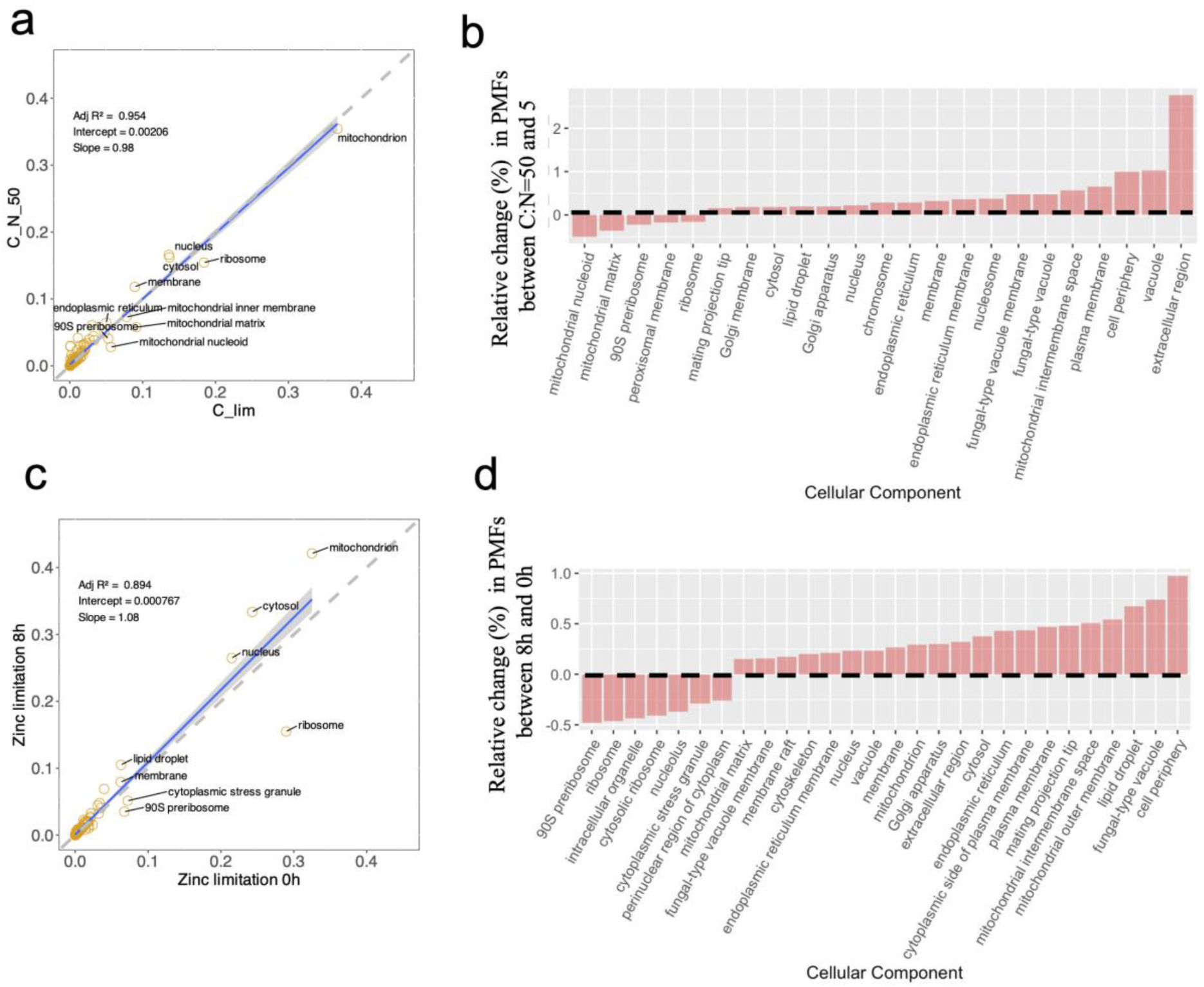
Protein resource redistribution at organelle level under nutrition limitation. Influence of N limitation on protein resource redistribution at organelle level. The proteomics samples were sampled from chemostat cultivation at C:N=5 (C limitation), 30, 50, 115, respectively (a). Relative changes in PMFs of organelles and sub-organelles under C limitation (C:N=5) and N limitation (C:N=50) (b). Influence of zinc deficiency on protein resource redistribution at organelle level. The proteomics samples were sampled at 0h, 4h, 8h, 12h, respectively, since the start of zinc deficiency (c). Relative changes in PMFs of organelles and sub-organelles between zinc deficiency at 0h and 8h (d). Here, only the cellular components with PMFs larger than 1%, of which the absolute relative change larger than 15% between two conditions, were displayed in graph b and d.

As another example, yeast could stop normal growth under serious zinc deficiency, thus providing chances to probe the resource allocation related to extreme nutrition limitation. In this aspect, the PMFs of organelles were recalculated using proteomics sampled at 0h, 4h, 8h, 12h since the zinc deficiency (24). Overall, PMFs of organelles at different time points were highly correlated (Fig.5b, adjusted R^2^ > 0.86). By comparison, as the extent of starvation increases, PMFs of organelles were more deviated from that at 0h. For example, using the linear fitting, the adjusted R^2^ at 12h is decreased to 0.876 compared to 0.958 at 0h. When looking into the deviation at the organelle level, it shows that lots of organelles’ PMF at 4h, 8h, 12h were significantly different from that at 0h, indicating the cell reallocates protein resources to respond to the zinc deficiency. Specially, we found that along with the zinc deficiency, the PMF of the ribosome decreased while the PMF of the mitochondria and cytosol increased, showing that protein for synthesis was reduced. Interestingly, it is noted that the PMFs of the lipid droplet and vacuole are increased accordingly, which is similar to the above nitrogen limitation (Fig. S11b). With zinc deficiency, the cellular resource would become limited, and the higher PMFs of the lipid droplet and vacuole may be related to enhanced catabolic activities within the cell, which could, to some extent, contribute to resource recycling and maintain normal cellular function.

Overall, it initially suggested that protein resources were relocated in a similar pattern under N limitation and zinc deficiency, and cells would resist the resource limitation by resource allocation at organelle levels.

### Organelle level resource allocation is genotype-dependent

Lastly, we further ask whether protein source allocation could be comparable among yeast species with different traits and genetic backgrounds. Here, using *I. orientalis* - a Crabtree-negative yeast species - as a typical example, we compared the PMFs of organelles between *S. cerevisiae* (Crabtree - positive) and *I. orientalis* (Crabtree-negative) under various growth conditions using a series of high-quality proteomic datasets generated by a recent study (16).

To probe the resource allocation at organelle levels in *I. orientalis*, we first assign component information to each protein from *I. orientalis* (Methods). Even with different procedures in protein location annotation, the numbers of proteins from different organelles follow a similar pattern to that of *S. cerevisiae*, with the nucleus having the most proteins, followed by mitochondria (Fig.6a). Three annotation procedures were utilized, and the number of proteins identified by ortholog relation is largest (Fig.6b). Once the protein location was annotated, the PMFs of organelles for *I. orientalis*, as well as the ρ of PMFs across samples, were calculated under batch, carbon-limited, nitrogen-limited, and phosphorus-limited growth conditions. In general, the resource allocation at the organelle level from *I. orientalis* could be comparable to that from *S. cerevisiae* (Fig.6c and 6d), lending evidence for the protein location annotation for *I. orientalis* used in this work.

**Figure 6.**
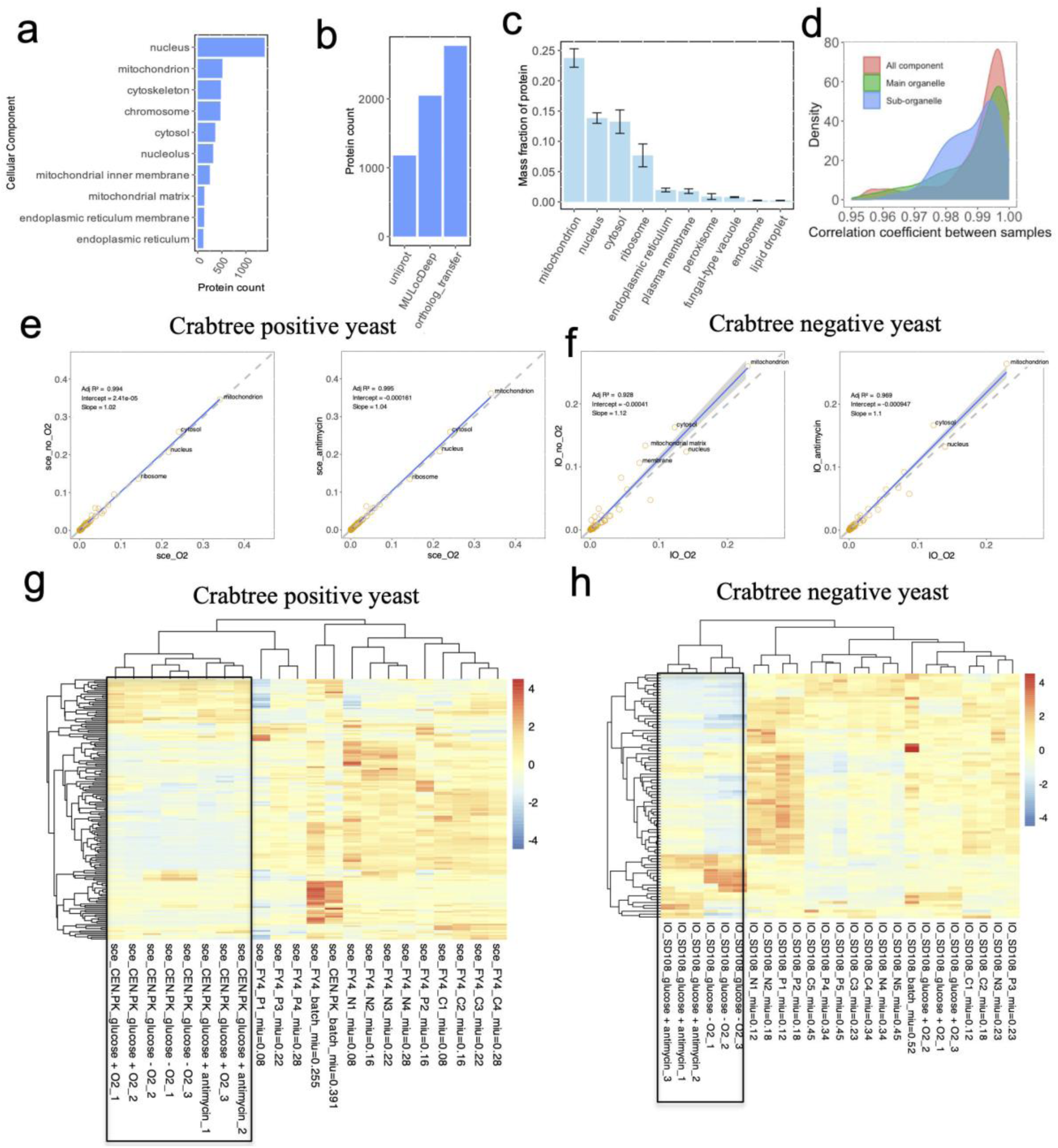
Comparison of resource allocation among organelles for two different yeast species - *Saccharomyces cerevisiae* (sce, Crabtree positive) *and Issatchenkia orientalis* (IO, Crabtree negative) under different conditions. Compartment annotation for IO (a). Numbers of proteins in compartment annotation from different sources for IO (b). Conservation and variation in mass fraction for proteins from main organelles of IO (c). Distribution of correlation coefficient calculated by protein mass fraction at organelle level across conditions for IO (d). Influence of the anaerobic and antimycin-inhibited condition on protein resource redistribution at organelle level for Crabtree positive and negative yeasts (e, f). Heatmap of mass fraction at organelle level across different conditions from the same study with cluster analysis of different samples for Crabtree positive and negative yeasts, in which the samples under anaerobic and antimycin-inhibited conditions from IO could be clearly separated from other conditions (g, h). Each row of heatmap represents a component.

Further, the PMFs of organelles under anaerobic conditions and respiration inhibition were compared to those under normal aerobic conditions for *S. cerevisiae* and *I. orientalis*, respectively. Obviously, we discovered that, when compared to *S. cerevisiae,* the oxygen limitation and respiration inhibition more significantly affect protein resource allocation at the organelle level for *I. orientalis,* partially consistent with the conclusion from (16). As shown in Fig.6e and 6f, it seems that protein resource allocation in *S. cerevisiae* remains relatively stable under these conditions, indicating greater robustness. In contrast, *I. orientalis exhibits* pronounced redistribution of protein resources. Specially, it could be found that the PMFs of the mitochondria and cytosol increase in *I. orientalis,* while the PMFs of the ribosome decrease under the above two conditions, likely due to the significant decrease in the growth of *I. orientalis* when respiration is repressed.

Further, we compared the PMFs of organelles for both yeast species under a series of different conditions and performed cluster analysis of samples based on those PMFs (Fig.6g and 6h). It is observed that for both yeast species, the batch cultivation could be distinguished from those chemostat cultivations under various nutrition (C/N/P) limitations. More importantly, it shows that, unlike *S. cerevisiae,* the cultivation of *I. orientalis* under O_2_ deprivation or respiration inhibition can be clearly distinguished from other conditions where the O_2_ is sufficient or the respiration is not inhibited, implying that the activity of oxidative respiration may play a significant role in regulating the protein resource allocation within Crabtree-negative yeast species like *I. orientalis*. Given the genomic differences between *I. orientalis* and *S. cerevisiae*, we hypothesize that this differential protein allocation is an adaptation to their specific ecological niches.

## Discussion

This study systematically explores how limited protein resource is reallocated among organelles within yeast cells under a wide range of conditions through leveraging the largest unified dataset of absolute quantitative proteomes (over 300 proteomes spanning 150 conditions) for two yeast species, *S.cerevisiae* and *I.orientalis*. The unit of protein abundance is important for proteome comparative analysis across conditions. Previously, the protein copy per yeast cell was calculated (11). However, the missed cellular size and density information will result in the bias during the unit transformation. In this work, we unified the protein abundance as mass fraction for reliable meta-analysis. In general, the organelle-level protein resource allocation was significantly affected by cell growth rate, nutrition limitation, and genome background. It initially shows that the growth law is still applied to the resource trade-offs at the organelle level, such as the protein resource being redistributed from major organelles, i.e., peroxisome and the nucleus, to ribosome to sustain fast growth. When the specific growth rate is controlled, the protein resource at the organelle level is redistributed for sustaining the cellular metabolism under extreme nutrition limitation. We also found that organelle-level resource allocation could be influenced by genotype. For example, the protein resource allocation is more robust in *S. cerevisiae* than that in *I. orientalis* under respiratory perturbations, reflecting that organelle-level resource allocation may be reshaped by long-term evolution. More importantly, for the first time, our integrated analysis shows that there exist close correlations between protein mass fraction, organelle level’s resource allocation and cellular physiological parameters. It suggests that the PMFs from a limited number of organelles could predict the cellular physiological parameters very well while a limited number of proteins could reflect the general resource allocation at the organelle level. It may verify that the organelle level resource redistribution was precisely tuned for cellular metabolic adaption at different scales. To make the most of the unified yeast proteomic dataset, an integrated online database was established for the community (https://yeast-proteome-database.streamlit.app), which provides curated proteomics data, enabling dynamic visualization and cross-condition comparisons of protein mass fractions (PMFs) at the organelle level. This comprehensive dataset will be further updated based on more new studies on yeast absolute proteomics analysis.

Our meta-analysis in yeast proteomics still faces several significant challenges, including batch effects and data heterogeneity, among which the variations in strain backgrounds, growth environments, and limit of detection by mass spectrometry further complicate cross-study comparisons, especially for low-abundance proteins. Note that the protein compartment annotations used in this work were mainly based on existing databases or computational inferences, which thus fail to capture dynamic relocalization of functional proteins under dynamic conditions (10), such as stress-induced protein migration between organelles (37). Additionally, the limitation in protein location annotation at the sub-organelle level makes it impossible to conduct accurate resource allocation analysis at a higher resolution. By addressing these challenges, future studies can reveal a more dynamic, high-resolution picture describing the resource allocation at multi-scale levels from major organelles to single proteins, which will help to profile the function of various organelles related to cellular fitness (38).

In summary, the unified large-scale absolute quantitative proteomic dataset, along with the online database, will serve as a valuable resource for yeast systems biology studies, enabling the exploration of the protein resource allocation at organelle levels under a wide range of varied growth conditions. By linking organelle-scale proteomics to cell physiology, metabolism, and evolution, this work sheds light on the principles in resource distribution for eukaryotes, setting a solid foundation to bridge the gaps between organelle functions and cellular metabolic adaptation.

## Methods

### Datasets collection for yeast

To collect all yeast absolute proteomic datasets, a thorough search of scientific literature in the past several years was conducted. Using platforms like PubMed and Google Scholar, keywords such as “*Saccharomyces cerevisiae*”, “yeast”, “proteomics”, “absolute proteomics” and “protein copy number” were employed to search the related studies. The proteome datasets were downloaded from each study. For each dataset, detailed documentation is crucial. A spreadsheet was used to record publication details (authors, title, journal, DOI), quantification method (e.g., SRM, iBAQ), and experimental conditions (growth medium, temperature, strain). Once all proteome datasets were downloaded, quality analysis was further conducted. The proteomes with the number of total measured proteins smaller than 1500 were removed. Meanwhile, the proteome, which cannot be converted into an absolute quantitative one based on the reference, were also filtered out. To address inconsistencies in protein names across different datasets, protein identifiers (ORF names, gene names) are standardized, often mapped to the standard locus name, facilitating the following data integration and analysis. In sum, the proteomes collected from (5, 7, 9, 11–13, 15–17, 24–27) (23, 28) were compiled and used for the following analysis.

### Unifying the unit of protein abundance across samples

To standardize protein abundance measurements from various studies into a common unit of mass fraction (f), which was defined as the mass of a specific protein divided by the total mass of all proteins within a cell.

For protein abundance in copies per cell (N_i_), obtain the molecular weight (M_i_) of each protein and calculate the weighted sum of all proteins (S) in the dataset.

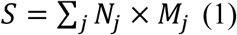

Where N_j_ the protein copy per cell and M_j_ the protein molecular weight. Next, the mass fraction of protein i (*f_i_*) is calculated by the following formula:

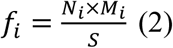

### Cluster analysis of samples based absolute proteomics

The process starts with data preprocessing, where the dataset is organized with samples as rows and proteins as columns. Preprocessing involves normalization—such as total protein normalization or log-transformation—to account for technical variability. Next, the package t-SNE(39) is applied to the pre-processed data, generating coordinates for each sample in the reduced space. Once computed, the results are visualized using scatter plots, where each point represents a sample.

### Compartment annotation for model yeast *S. cerevisiae*

To annotate the compartment for all proteins in *S. cerevisiae*, the professional *Saccharomyces* Genome Database (SGD) at https://www.yeastgenome.org/ was utilized (34). The Gene Ontology (GO) annotations were downloaded for each protein from SGD. Next, based on GO annotation, the “Cellular Component” section to identify the cellular compartments where the protein is located was extracted. SGD provides reliable and current annotations, curated from literature and high-throughput experimental studies, ensuring a comprehensive understanding of the protein’s cellular location. Thus, based on the SGD annotation, the initial mapping between compartment and protein was established for most proteins. Moreover, for proteins from several widely studied organelles-mitochondria, nucleus, cytoplasm, nucleolus, fungal-type vacuole membrane, fungal-type cell wall, and plasma membrane - the manual annotation recorded from SGD was mainly used for all the analysis in this work. However, by comparison, it displays differences in the sub-organelle annotation between literature and SGD for proteins from mitochondria. Thus, in these specific cases, the experimental protein location will be used to update the original annotation for proteins at different sub-organelles from mitochondria recorded in SGD. Otherwise, the protein compartment annotation from other organelles or sub-organelles recorded in SGD was utilized.

### Compartment annotation for non-model yeast

To annotate the subcellular compartments of proteins in non-model yeast species, three complementary methods — UniProt-based annotation (40), ortholog-based annotation, and deep learning-based prediction — were employed, respectively. Firstly, if a protein for non-model yeast has existing compartment annotation from UniProt, the recording will be kept. For the protein without annotation from UniProt, the ortholog-based annotation was used. In detail, the evolutionary relationships between the non-model yeast proteins and those of *S. cerevisiae* were identified. The toolbox (41) was employed to establish the reciprocal best hits between the two species, allowing the transfer of known subcellular localization data from *S. cerevisiae* (sourced from databases like SGD) to the non-model yeast proteins. Lastly, for the remaining proteins without compartment annotation, the deep learning-based prediction using MULocDeep (42) to predict compartments from protein sequences in FASTA format was utilized. For the output from MULocDeep, the high-confidence predictions are favored, while those with low scores are considered less reliable. Finally, the annotations from all three methods were unified and merged to represent the final protein compartment for non-model yeast.

### Calculation of volume and surface area for each protein of yeast

There are two procedures in this work for calculating the protein volume. In the first procedure, the toolbox named ProteinVolume v1.3 (43) was used. For each protein structure, the Van der Waals Volume and Void Volume were calculated by ProteinVolume v1.3 respectively. Then the total volume of a protein structure was found as the sum of Van der Waals Volume and Void Volume. In the second procedure, the specific volume of a protein is calculated based on the assumption that all proteins have the same density, about 1.37 g/cm^3^. Assuming the protein has the simplest shape, a sphere, the radius is calculated based on the following formula (44):

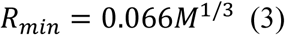

Where M is the molecular weight of a protein (the unit is Dalton), and R_min_ is the radius (the unit is nanometer).

Once the R_min_ was calculated based on formula 1, the protein structure volume was further calculated using formula 2.

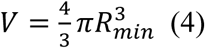

Where V is the volume of a protein structure.

### Calculation of protein mass fraction at organelle level

Once all protein abundances were changed to mass fractions, the protein mass fractions at the organelle level could be calculated. For each organelle, its encompassed proteins were collected, then the mass fractions of all proteins within the organelle were summed together to get the total mass fraction of proteins from each organelle. It should be noted that one protein mass fraction may be repeatedly calculated for some organelles, as the protein could occur in multiple organelles based on the compartment annotation from the SGD database.

### Construction of an online database for yeast absolute proteomics

For the easy usage and visualization of yeast proteomic datasets collected in this work, an online database was built. A unique ID (like P1, P2) was given for each sample for the online version of proteomics. Then the detailed condition was assigned to the above unique ID. The on-line database thus includes four types of datasets: the PMF across conditions, the PMF at the organelle level across conditions, the mapping between proteins and organelles (which is mainly downloaded from SGD) and the environmental condition for each sample. The online database also provides some functions for the quick analysis and visualization of proteomics, as shown in this study. For the actual analysis, the users could refer to the introduction of our online database (http://prdtst.tsynbio.com:51443/database/).

### Establishment of formulas between organelle level PMF and yeast phenotypic datasets

A generalized linear model was constructed to quantify the relationship between organelle-level PMF and yeast phenotypic data. The model accommodates one or more input features and follows the similar formula:

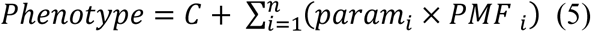

Where C is the intercept term, param*_i_* represents the regression coefficient for the *i*-th organelle’s PMF, and PMF_i_ represents the PMF for organelle *i*.

### Correlation analysis between protein mass fraction and organelle level mass fraction

To identify a minimal predictive protein set for organelle-level resource allocation (specifically the 24 major organelles), we employed recursive feature elimination (RFE) coupled with ridge regression (10-60 proteins, step size=1). Ridge regression was selected as it demonstrated superior performance (highest R²) in preliminary full-feature modeling. The dataset was split with 20% reserved for testing. During each RFE iteration, features were recursively eliminated based on ridge regression coefficients, retaining only the top n most predictive proteins at each step (where n decreases from 60 to 10). Model performance was evaluated using R² and RMSE on the test set. Subsequent KEGG pathway enrichment analysis of the minimal protein set was performed using the clusterProfiler R package.

### Protein-protein interaction (PPI) network analysis

The protein-protein interaction (PPI) network was constructed utilizing the STRING database (version 12.0) (36), followed by k-means clustering and topological characterization.

## Quantification and Statistical Analysis

For two group comparisons in this work, a two-tailed Wilcoxon rank sum test was calculated.

## Data availability

More detailed results in this study are available on https://yeast-proteome-database.streamlit.app/ and https://github.com/hongzhonglu/large_scale_yeast_proteomics_analysis.

## Acknowledgments

This work is supported by grant 2022YFA0913000 from the National Key R&D Program of China, grants 22208211 and 22378263 from the National Natural Science Foundation of China (NSFC). We thank the help from Hao Yu and Xueting Wang in the data analysis.

## Author Contributions

H.L. and J.N. designed the project. H.L. performed the research and wrote the manuscript. YZ built the on-line database and ZZ conducted the formula fitting analysis. All authors reviewed and edited the manuscript. H.L obtained the funding for this study.

## Supplementary figures

**Supplementary Figure 1.**
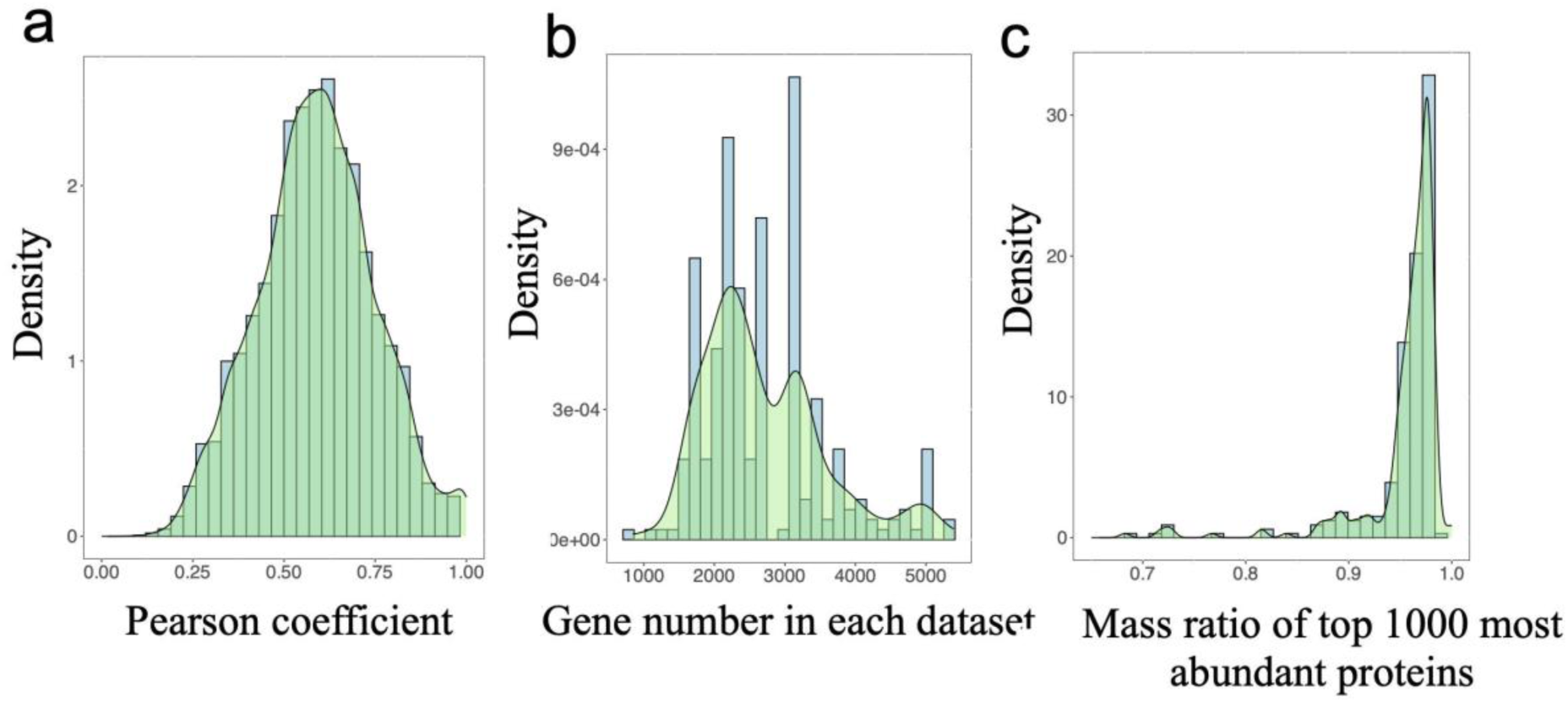
Quality analysis of protein abundance. Pearson correlation coefficient in mass fraction of organelles across different samples (a). Density of protein (gene) number across all datasets (b). Density of mass ratio from top 1000 most abundant proteins (c).

**Supplementary Figure 2.**
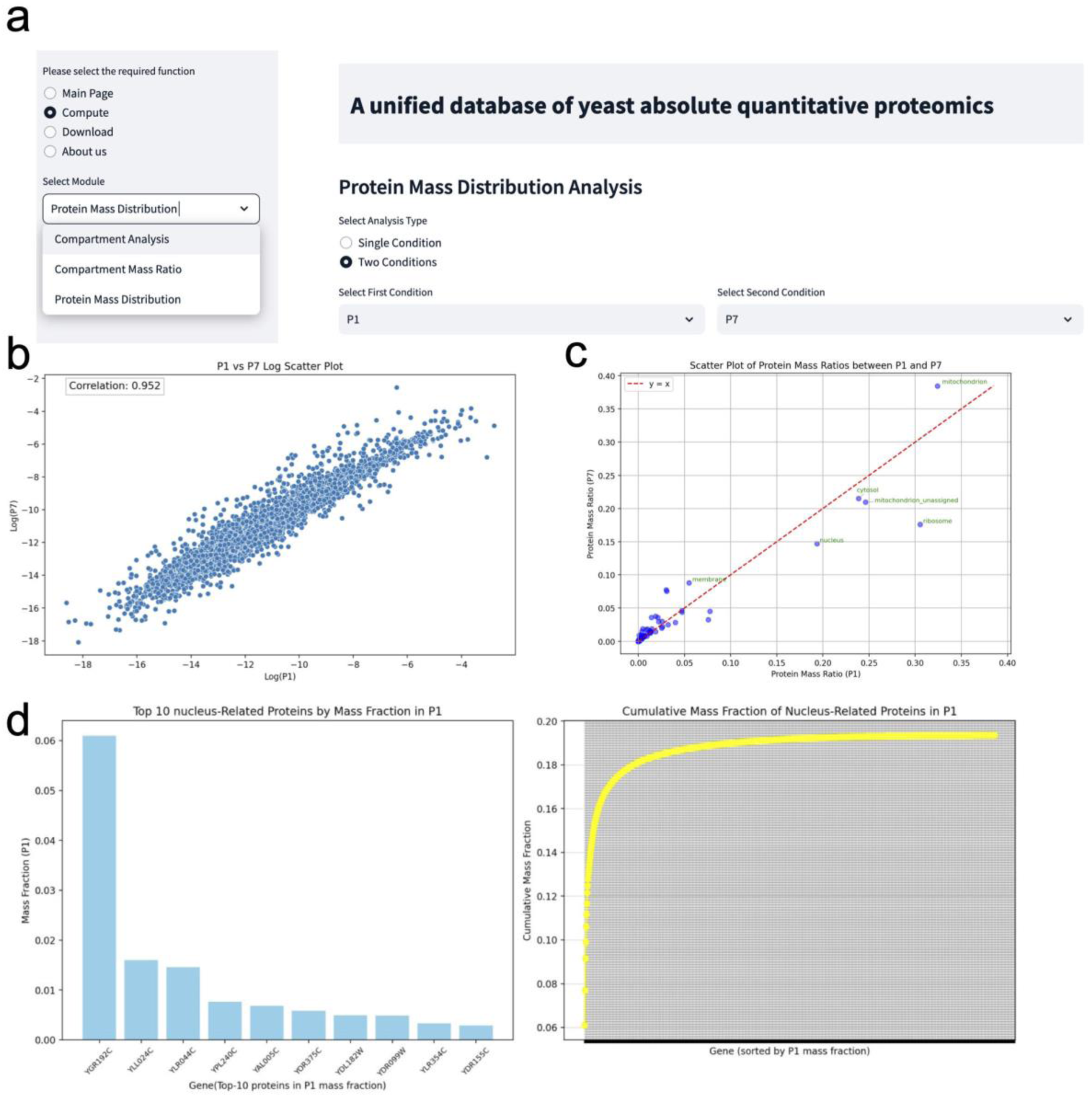
Screenshot of online database for visualization of yeast absolute quantitative proteomics. Main functional modules of the online database (a). Correlation analysis based on protein mass fraction between two conditions (b). Correlation analysis of protein mass fraction at organelle level between two conditions (c). Rank analysis of protein mass fraction in specific organelle (d). P1 and P7 represents proteomic datasets from different conditions or studies.

**Supplementary Figure 3.**
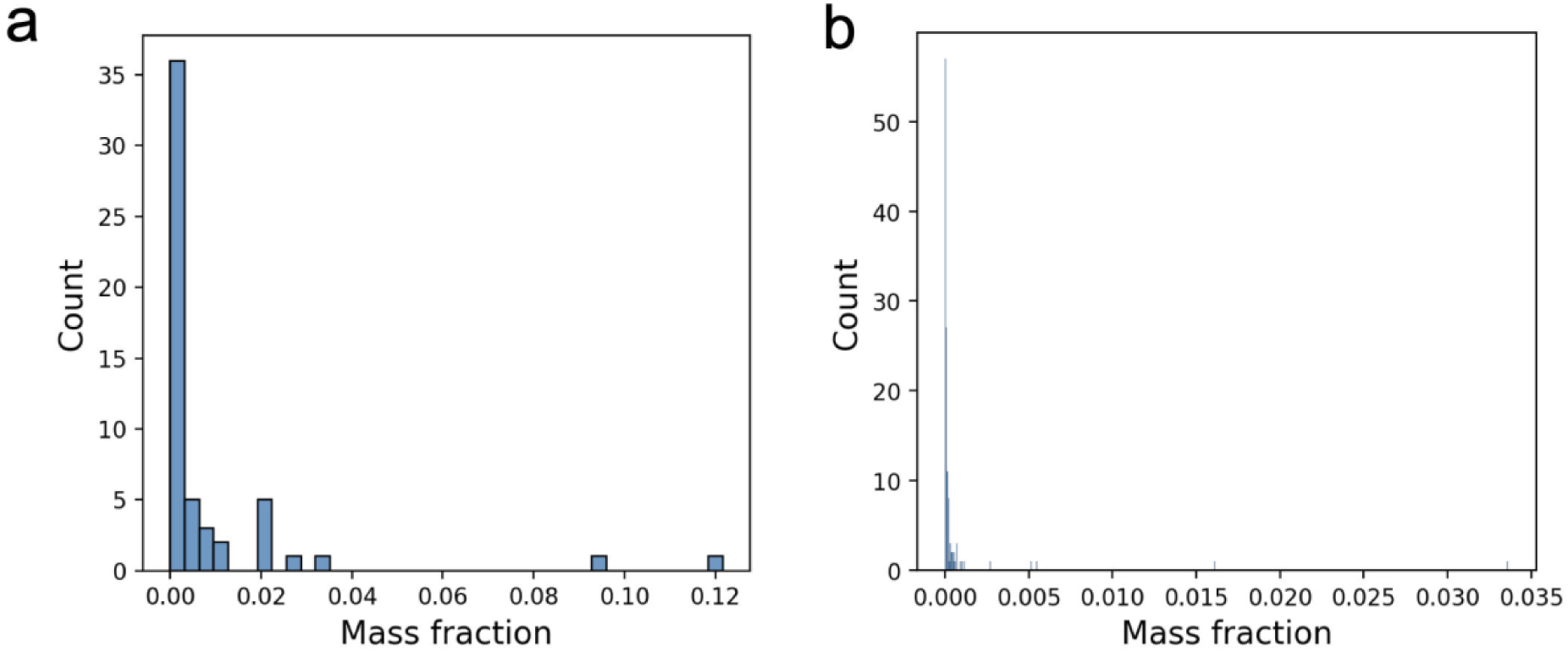
The percentage of proteins shared by two organelles per total protein based on the annotation from SGD database (a). The percentage of proteins which could transfer across two organelles under different cell cycles per total protein (b).

**Supplementary Figure 4.**
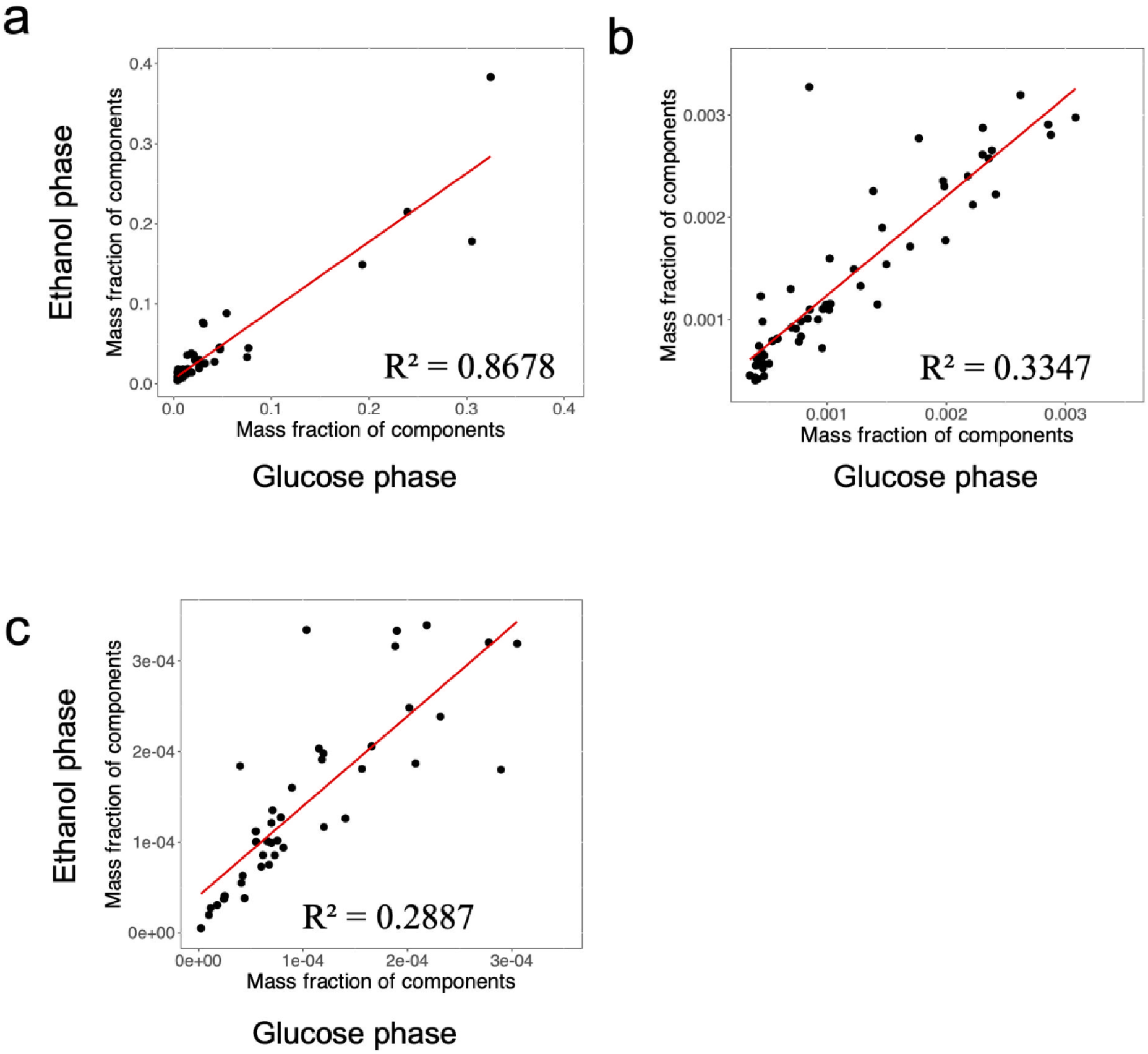
Protein distribution is more robust for cellular components with larger protein mass fraction. Here two different growth conditions were selected for the correlation analysis. Correlation of protein mass fraction from large component with protein mass fraction ≥ 0.0035 (a). Correlation of protein mass fraction from medium component with protein mass fraction in range of 0.00035 to 0.0035. (b). Correlation of protein mass fraction from small component with protein mass fraction < 0.00035 (c). The proteomic datasets in batch cultivation under glucose and ethanol phases were from www.pnas.org/cgi/doi/10.1073/pnas.1918216117.

**Supplementary Figure 5.**
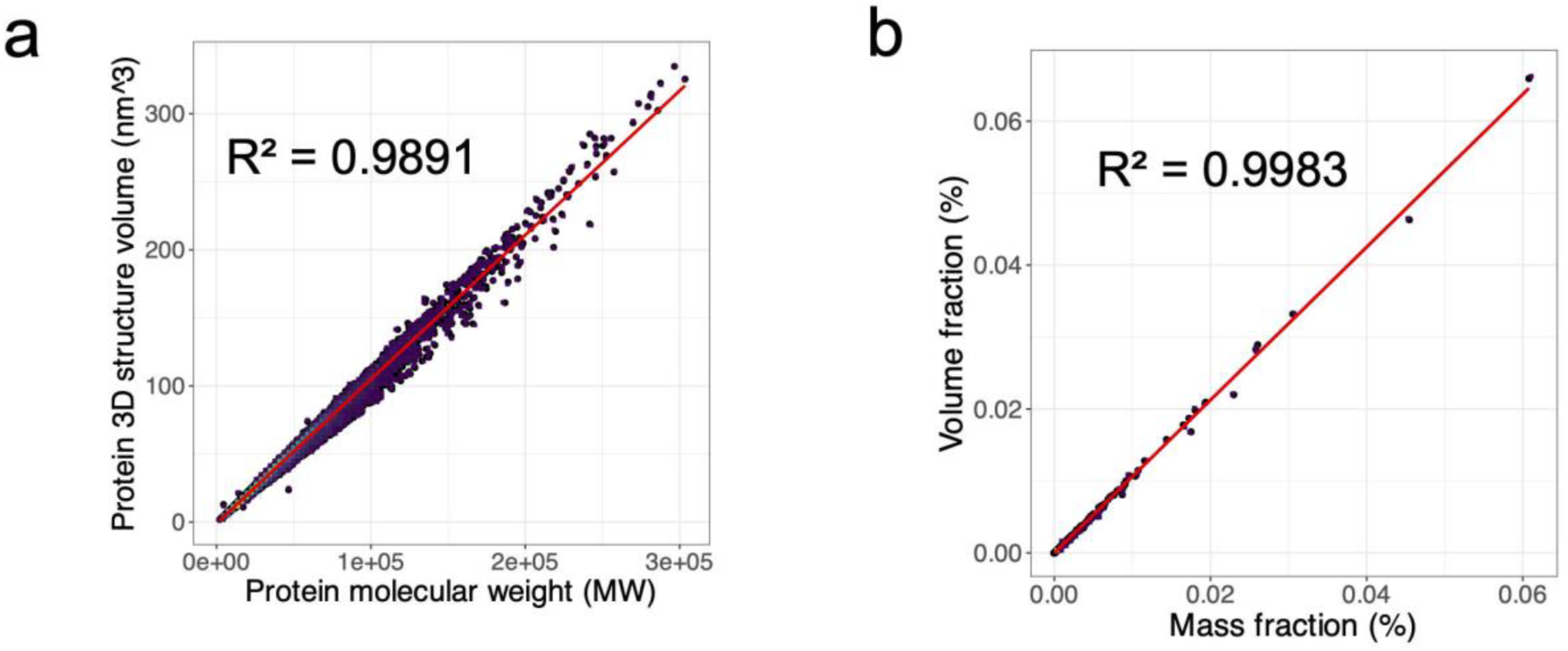
Linear correlation exists between volume and mass for single protein from *S.cerevisiae* (a). Linear correlation exists between mass fraction and volume fraction for proteins from *S.cerevisiae* under one batch condition when considering the protein abundance (b).

**Supplementary Figure 6.**
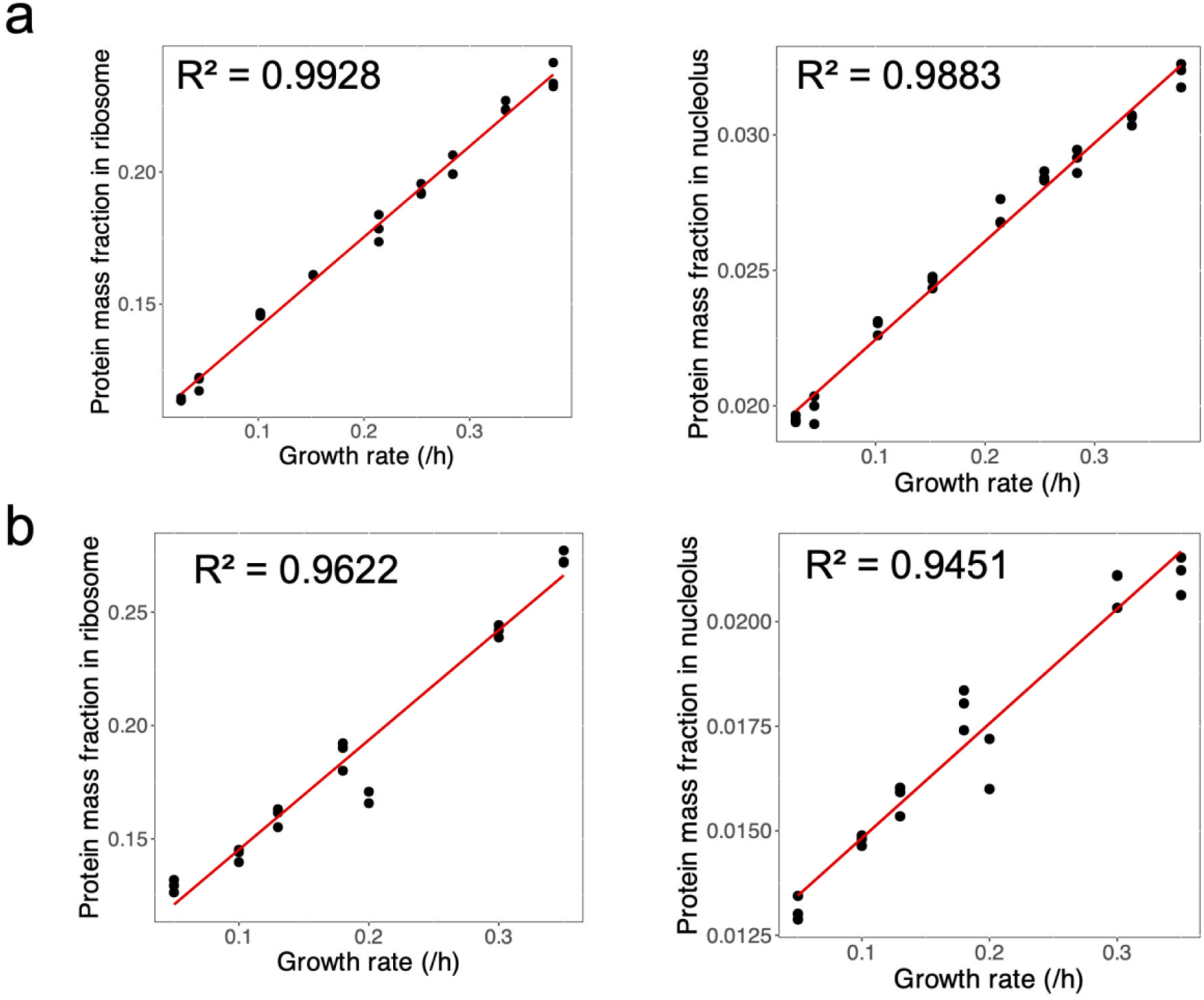
Mass fractions of proteins from nucleolus and ribosome are linearly correlated to growth rate under carbon-limited (a) and nitrogen limited conditions (b).

**Supplementary Figure 7.**
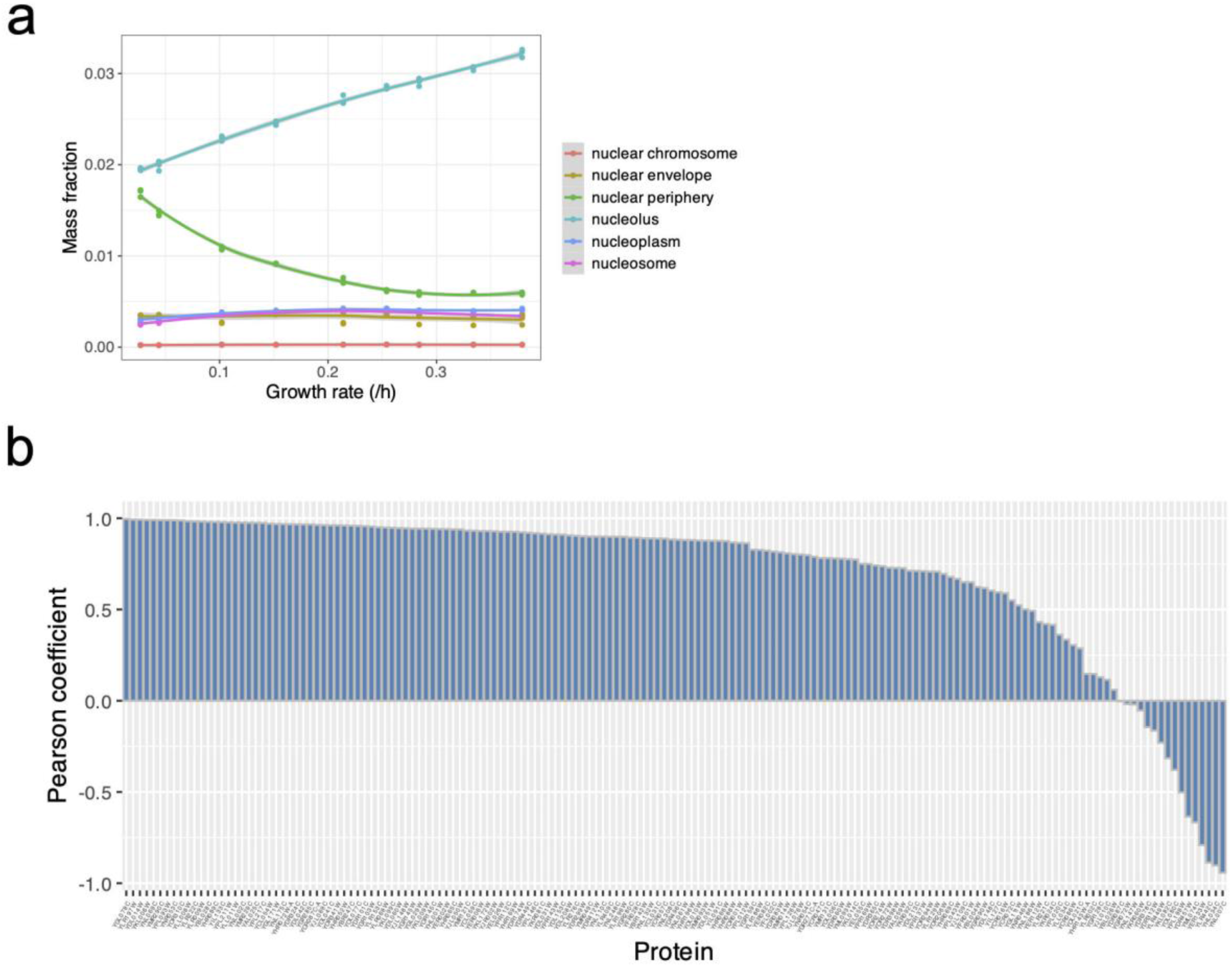
Protein resource allocation within nuclear is influenced by the growth rate. Tendencies in mass fraction of proteins from major components of nuclear along with growth rate (a). Correlation of growth rate and mass fraction for single protein from nucleolus (b).

**Supplementary Figure 8.**
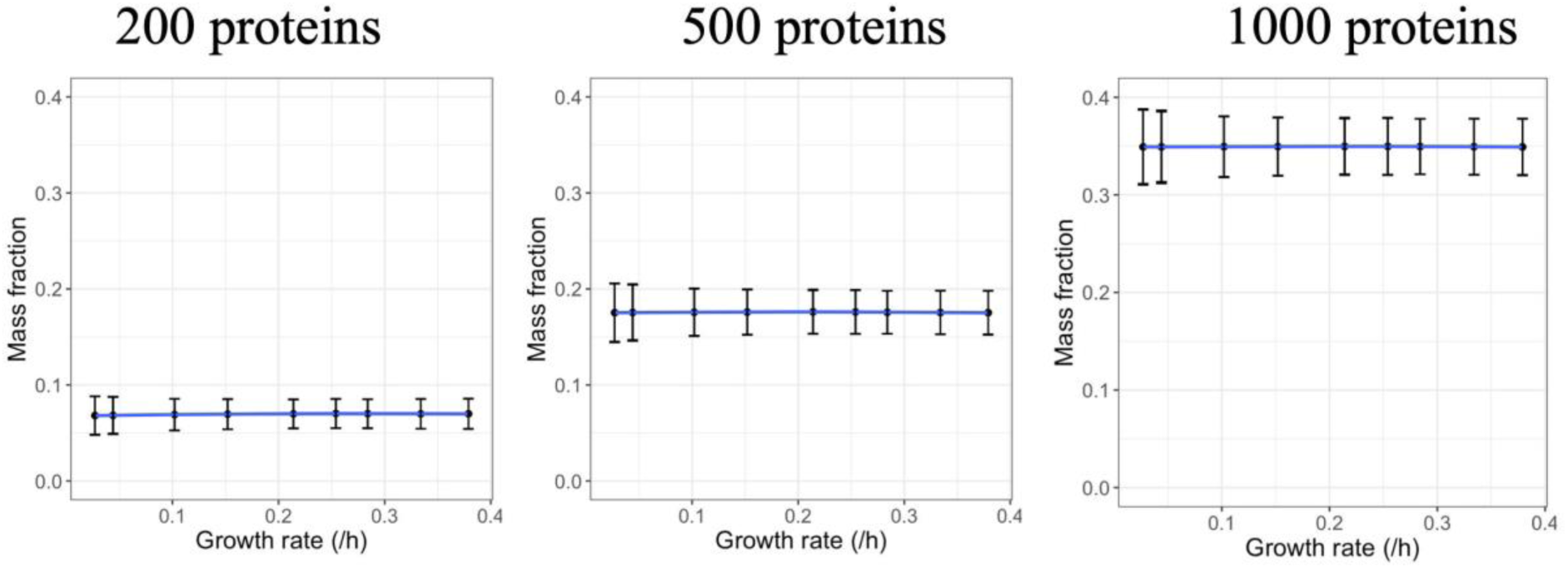
Mass fraction of randomly selected 200, 500 and 1000 proteins under different growth rate. For each test, the proteins were randomly sampled 1000 times to calculate the average and SD values at each growth rate.

**Supplementary Figure 9.**
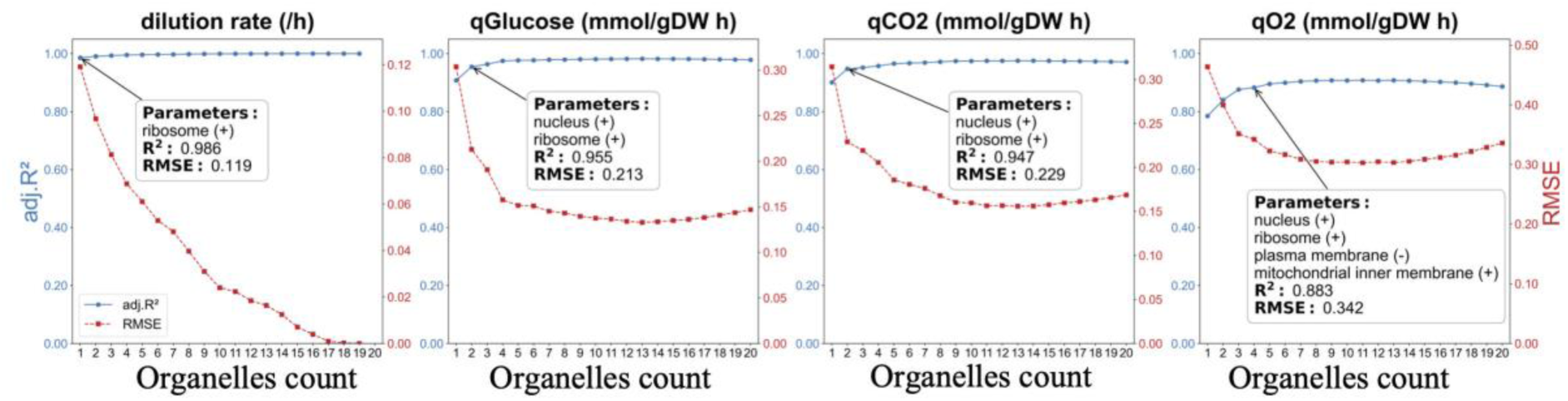
By increasing the number of organelles in the formula fitting for prediction of yeast physiological parameters, the R^2^ can be appropriately increased, and the RMSE can be reduced accordingly. The arrow represents the inflection point in each profile.

**Supplementary Figure 10.**
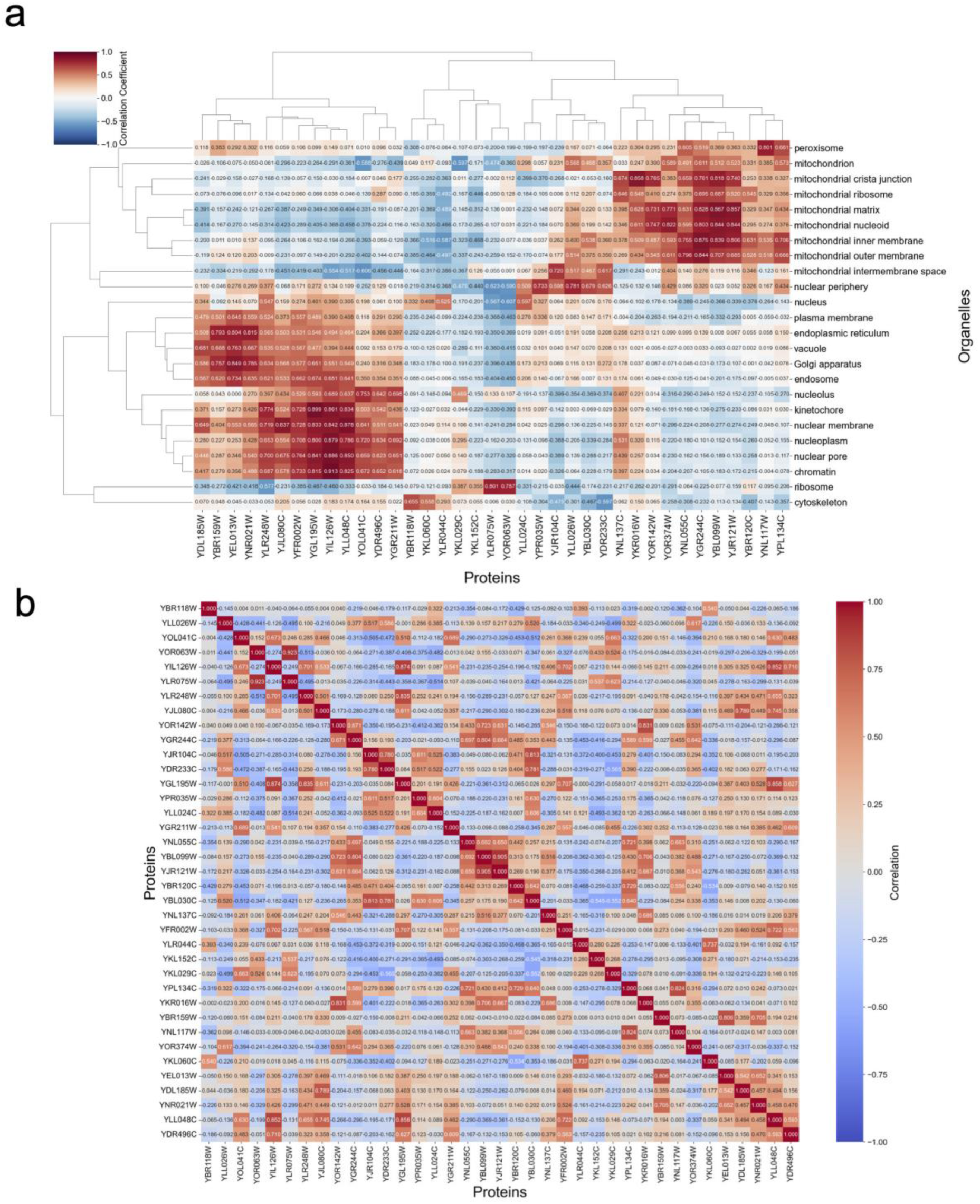
Correlations between the mass fraction of 37 proteins and the mass fraction of proteins within 24 organelles and sub-organelles (a). Correlations among 37 proteins in the mass fraction (b).

**Supplementary Figure 11.**
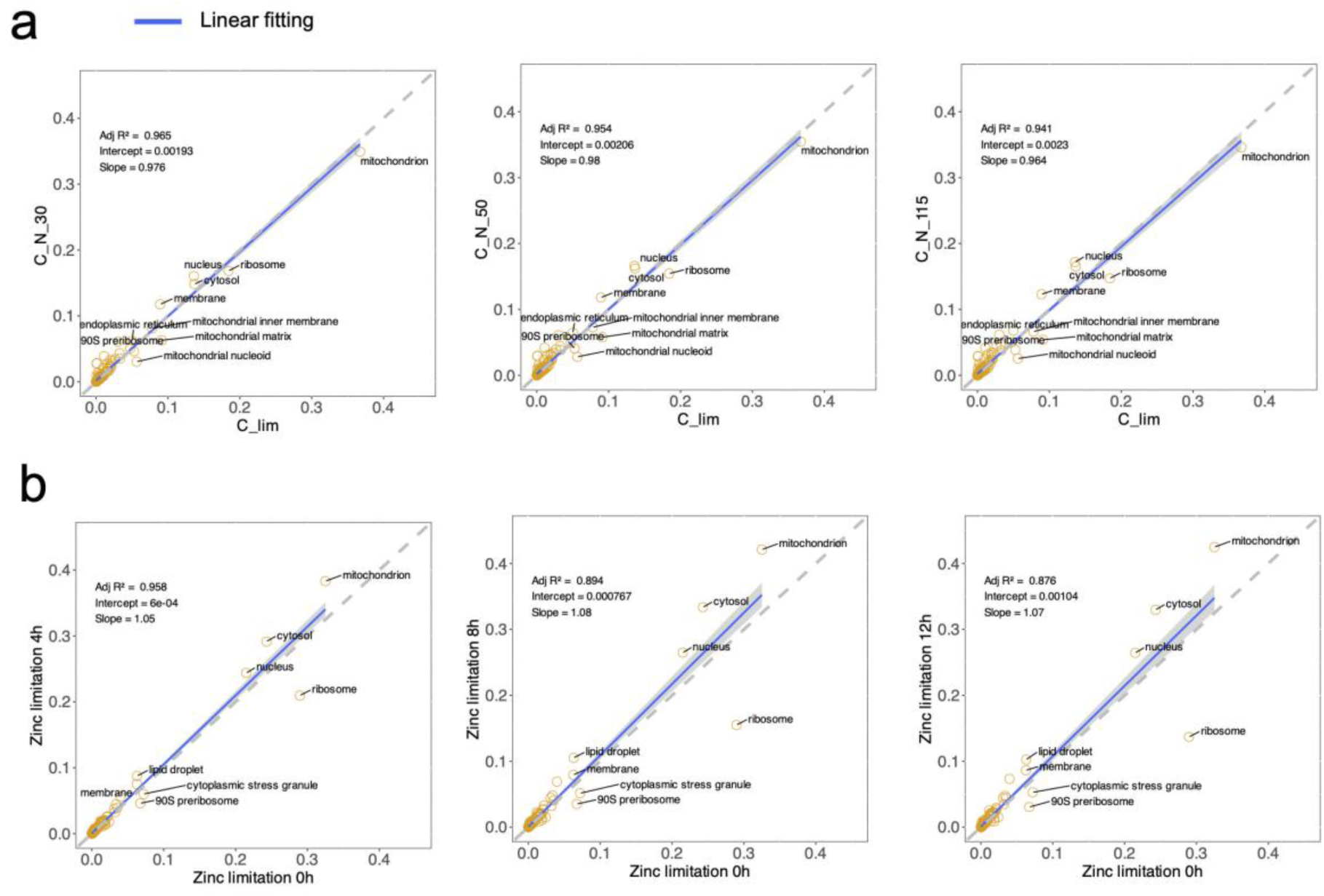
Relative change in mass fraction of proteins at cellular component level under nutrition limitation. Influence of N limitation on protein resource redistribution at organelle level. The proteomics samples were sampled from chemostat cultivation at C:N=5 (C limitation), 30, 50, 115, respectively (a). Influence of zinc deficiency on protein resource redistribution at organelle level. The proteomics samples were sampled at 0h, 4h, 8h, 12h, respectively, since the start of zinc deficiency (b).

